# Consequences of phenological shifts are determined by the number of generations per season

**DOI:** 10.1101/2025.02.17.638696

**Authors:** Heng-Xing Zou, Volker H.W. Rudolf

## Abstract

The order and timing of species’ arrival in a community can often change species interactions. Termed priority effects, this phenomenon arises in communities with drastically different life histories (e.g., bacteria, insects, and amphibians), yet how such life history traits affect consequences of priority effects in seasonal systems remains understudied. Here, we test how a key aspect of life history, numbers of generations in a growing season, interact with priority effects using two competing flour beetles, *Tribolium castaneum* and *Tribolium confusum*. We manipulated the sequence and timing of arrivals and collected all adults after one, two, or three generations, simulating phenological shifts in communities with different growing season lengths. The early species always reached a higher population, and in general, its per-capita competition towards the late arriver was stronger, indicating priority effects. However, fitting competition models to the data revealed that more generations in a season led to smaller ranges of the calculated stabilization potentials and fitness differences between species and more positive stabilization potentials, indicating that overlapping generations buffered interactions against strong priority effects from different arrival times. These results suggest that effects of phenological shifts (e.g. due to climate change) on species coexistence mechanisms are contingent on the life-histories of interacting species.

## Introduction

The relative timing of seasonal life history events (phenology) can often change species interactions. Among competing species, this phenological difference can lead to priority effects, affecting species coexistence and long-term community dynamics (Fukami 2015; Rudolf 2019; Zou and Rudolf 2023). Despite extensive studies, priority effects among systems with different life history traits, such as the length of the life cycle and seasonality, are rarely compared and synthesized (Zou and Rudolf 2023). These biological differences make the development of a unified theory of priority effects difficult because certain mechanisms and consequences may only apply to specific systems.

Priority effects have been detected in microbial communities to plants, insects, and vertebrates (Rudolf 2018; Toju et al. 2018; Blackford et al. 2020; Holditch and Smith 2020). The differences between these taxa might seem too large to warrant any meaningful comparison, yet they all share a key biological process: reproduction in a periodic environment. Nectar microbial communities assemble when their hosts flower, and annual plant communities complete their life cycle within a growing season. This periodic (seasonal) reassembly of communities not only provides opportunities for priority effects but also evolution of different reproduction regimes.

For instance, in an ephemeral pond that has a wet period (growing season) of several months, the amphibians and fish communities may only reproduce once, while the zooplankton and protist communities may produce multiple generations. Evaluating the effect of such differences in reproduction regimes on priority effects is inherently difficult between distantly related taxa, such as the amphibian and protist communities in the above example. Nevertheless, traits such as the number of reproductive events can differ between closely related taxa or display surprising phenotypic plasticity within a species in response to changing environments. For instance, the development rate of *Daphnia* is higher in urban areas with higher temperatures, leading to a faster “pace of life” that enables more frequent reproduction (Brans and De Meester 2018). Persistent warming increases the number of generations in a year in many European lepidopterans (Altermatt 2010). These examples provide opportunities for evaluating the interaction between strengths of priority effects and life history traits, specifically the number of generations in a growing season.

Theory predicts that such differences or changes in the number of generations in a growing season can alter the strength of priority effects among species, especially when the priority effect arises from the difference in size or developmental stage of the initial generation (i.e., trait-dependent priority effects; Zou and Rudolf 2023; Zou et al. 2023). In these communities, the early species grows to a larger size or later stage when the late species arrives (colonizes or emerges from dormancy), leading to a competitive advantage that results in priority effects (Shorrocks and Bingley 1994; Connolly and Muko 2003; Rasmussen et al. 2014; Blackford et al. 2020). With more generations in a growing season, continuous reproduction leads to overlapping generations, and the populations of both species consist of multiple coexisting stages/sizes. As a season contains more overlapping generations, the size/stage distribution of the two species becomes less different, lowering the importance of the initial size/stage difference and weakening trait-dependent priority effects (Zou et al. 2023). However, this prediction lacks empirical evidence and is solely based on priority effects generated by size/stage difference, while in nature, they could also arise from other mechanisms. For instance, arriving early may lead to a higher abundance, and the early arriver advantage may be maintained through positive frequency dependence that does not change by the number of generations (i.e., frequency-dependent priority effects; Ke and Letten 2018; Toju et al. 2018; Zou and Rudolf 2023).

Alternatively, differences in arrival time can allow early arriving species to consume smaller later arriving species (intraguild predation; Rasmussen et al. 2014), and if the predation rate is high, the late species may not survive to adulthood, leading to consistent exclusion regardless of the number of generations.

To test this prediction and gain more insight into the interaction between the number of reproductive events and phenological shifts in nature, we conducted an experiment with two competing flour beetles, *Tribolium castaneum* (red flour beetles) and *Tribolium confusum* (confused flour beetles). We experimentally created an arrival time gradient, then allowed the beetles to grow and reproduce for different durations to simulate different numbers of overlapping generations in a “growing season”. We show that the two species competed strongly and displayed priority effects, but the effect of the number of generations on priority effects is more nuanced than previous theoretical predictions.

## Methods

### Study Organisms

*Tribolium castaneum* and *Tribolium confusum* have been widely used as a model system to study interspecific competition for a century (Park 1948; Edmunds et al. 2003; Pointer et al. 2021). They feed on flour and grains, compete readily in the same habitat (Park 1954), and display priority effects (Holditch and Smith 2020). We used a black strain of *T. castaneum* collected from Hereford, TX (J.P. Demuth) and strain PAK of *T. confusum* purchased from the USDA Agricultural Research Service (Manhattan, KS). Life history traits of the two species differ by their strains; in our experiment, *T. castaneum* develops faster than *T. confusum* (Park 1957), but the latter produces more eggs under our experimental conditions (Table S2). Beetles were reared in incubators at 32°C and 20-40% relative humidity with constant airflow. To minimize the effects of pathogens and parasites, we washed eggs collected from stock culture with 10% bleach solution for 30s to 1min, then incubated the eggs in flour (King Arthur, Norwich, VT) with 0.03% fumagillin (KBNP Inc., Anyang, South Korea) by weight. We conducted all experiments by adding eggs to 7-dram vials (29mm diameter, 50mm height; Bel-Art, Warminster, PA) with perforated caps to allow for air exchange. Pilot trials showed that under the experimental densities, 8g medium per vial induces measurable competition. To lower the potential effects of the maternal environment, we collected eggs from newly emerged adults reared under the same environmental conditions. We collected eggs once per three days to maximize synchrony among larvae while maintaining the large number of eggs needed.

### Experimental Design

We conducted a full factorial design of seven relative arrival times, two total densities, three ratios between *T. castaneum* and *T. confusum*, and three season lengths. To simulate the difference in arrival times, we added eggs of one species at day 0, then added eggs of the competitor either on the same day (simultaneous arrival) or 6, 12, or 18 days later, yielding seven temporal treatments (Figure 1A). We define relative arrival times as the arrival time of *T. castaneum* minus that of *T. confusum*, such that a negative value indicates the early arrival of *T. castaneum* and vice versa. For each temporal treatment, we used a response surface design (Inouye 2001) with two total densities (30, 60) and three density ratios (1:2, 1:1, 2:1, in the order of *T. castaneum* : *T. confusum*) to measure the per-capita interspecific competition (Figure 1B), resulting in a total of 126 competition treatment scenarios.

**Figure 1.**
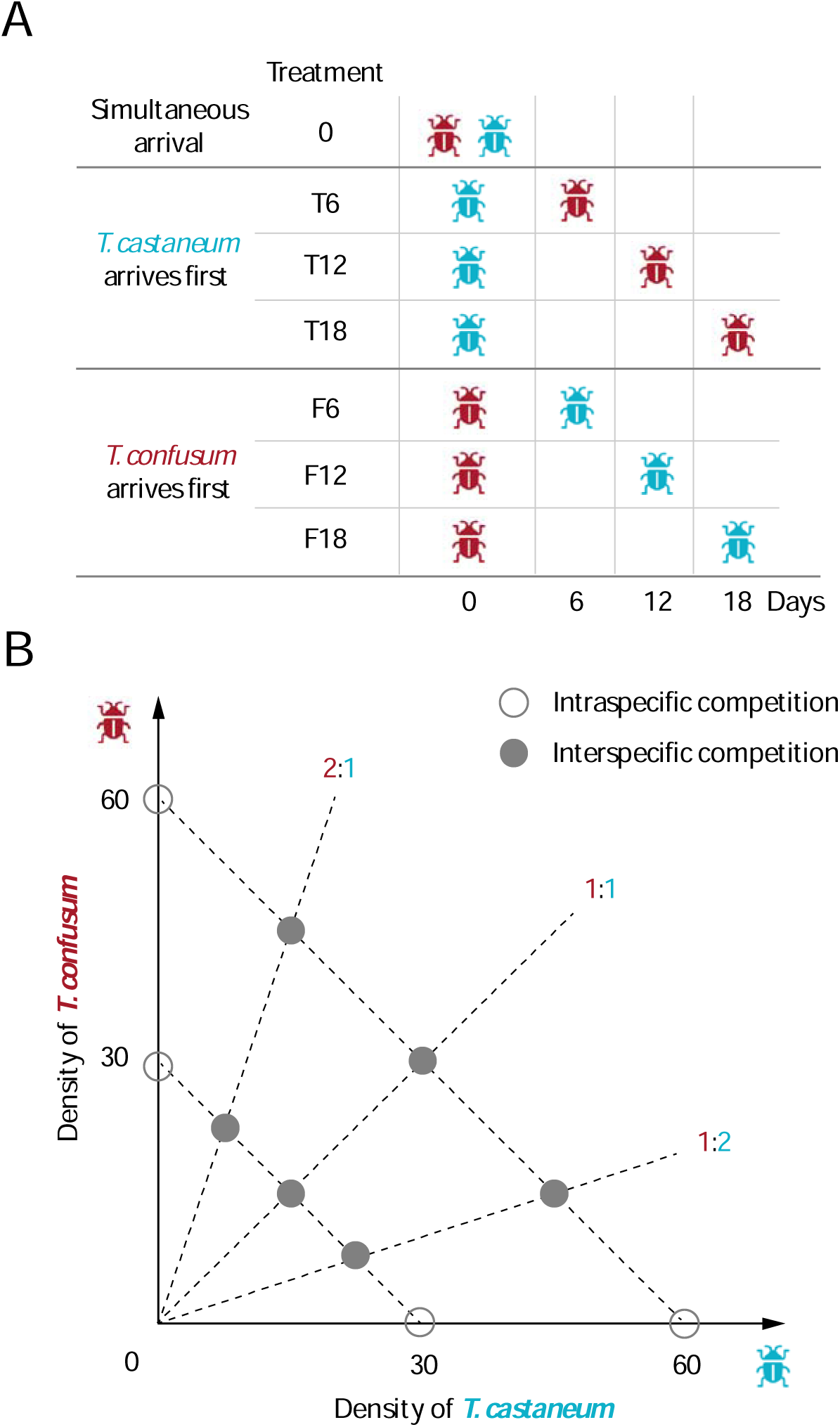
Scheme of the experiment. A. Different arrival times (temporal treatments) of the two species, *Tribolium castaneum* and *T. confusum*. B. Response surface design with six density combinations at two total densities and three ratios, along with two single-species treatments, to evaluate competition coefficients. Each treatment in the response surface is replicated three times.

In addition, we set up single-species vials with 30 or 60 eggs to evaluate the intrinsic growth rates and intraspecific competition; these vials were not multiplied by temporal treatments. Finally, we simulated different numbers of generations in a season by growing the vials for 36, 72, and 108 days, corresponding to one, two, and three generation times per season. We determined the length of each season based on the 36 days from eggs to adults for both species under our experimental conditions. With three repetitions per single and two species treatment combinations, this factorial design yielded 390 total experimental units. We started the experiment on May 21, 2023.

To quantify population sizes, we started counting adults at the assigned season length after adding the first species (day 36, 72, or 108). We repeated this process every six days thereafter until all live individuals emerged as adults, at which time we terminated the vial. At each census, we kept all adults in the vials to maintain the competitive pressure as other individuals developed but removed eggs to terminate further reproduction. This allowed the larvae and pupae to complete their life cycle and develop into adults, at which stage we can easily identify them to species (while juvenile stages look identical). This also ensured that arriving early would not lead to a longer time for population growth because lengths of seasons were fixed, and eggs were removed. We changed the medium of all experimental vials every 30-40 days. If the change date did not coincide with a check date, we retained all individuals, including eggs, while also recording the number of adults. The experiment ended on Oct 27, 2023.

### Statistical Analysis

To detect priority effects, we first fitted linear mixed effects models for each species, with the final adult population as the response variable and relative arrival time, the number of generations, and the ratio of initial egg densities as predictor variables and vial ID within each treatment as the random effect. The ratio of initial egg densities was fitted as a categorical variable. We fitted the full model separately for each species using the R package lmerTest (Kuznetsova et al. 2017), then selected the best model using the step() function in the lmerTest package.

To further quantify the competition between flour beetles, we parameterized discrete-time Lotka-Volterra models for vials from each arrival time and the number of generations under a Bayesian framework. Fitting Beverton-Holt and Ricker models either led to issues with convergence (>1000 divergent transitions after warmup) or lower mean log likelihood (see Supporting Information, Figure S1-S2 for details). Key traits of flour beetles, including fecundity, are highly dependent on the natal habitat (Van Allen and Rudolf 2013, 2013). Therefore, we cannot estimate the intrinsic growth rate of each species by measuring the fecundity of a single individual in fresh media because its natal habitat would be different from those in vials with multiple generations per season. Instead, we used information from our single-species vials to estimate intrinsic growth rates at different generations per season and then used this information to assist in the estimation of parameters in the two-species model.

First, we fitted single-species Lotka-Volterra models to data obtained from single-species vials of each number of generations in a season:

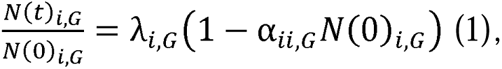

Where *N*(*t*)*_i,G_* is the final adult density of species *i* with *G* generations, *N*(0)*_i,G_* is the initial egg density, and α*_ii,G_* is the per-capita intraspecific competition coefficient. In this step, we used flat priors for λ*_i,G_* and α*_ii,G_* but confined the latter from 0 to 1. We then used the fitted intrinsic growth rates to inform the model fitting of two-species Lotka-Volterra models:

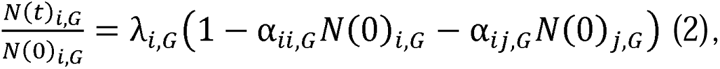

where α*_ij,G_* is the interspecific competition coefficient from species *j* to species *i*. We constructed a normal distribution with mean as the posterior estimates of each λ*_i,G_*. To assist convergence, we constrained the standard deviation as 10% of the mean λ*_i,G_*. We repeated the above process, fixing the priors of λ*_i,G_*for all vials with *G* generations, but used flat priors for the competition coefficients α*_ii,G_* and α*_ijG_* from 0 to 1 to allow for their variations at different relative arrival times. This approach assumes that the intrinsic growth rate changes with the number of generations but not relative arrival times. This is the most likely biological scenario because the number of generations is naturally linked with the reproductive outcome, while any effect of relative arrival times on the maternal age and the environment will be reflected by competition coefficients.

We randomly subsampled 1,000 posterior estimates of each competition coefficient from the model fitting result and calculated stabilization potential (SP) and relative fitness differences (RFD) from each sampled group of the four coefficients (Ke and Letten 2018; Godwin et al.

2020); these metrics are often used to predict outcomes of competition by comparing the two species’ self-limitation vs. limitation on the other species. We calculated SP and RFD as:

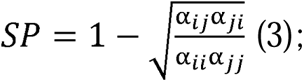

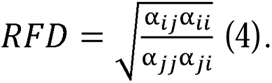

The two species coexist if 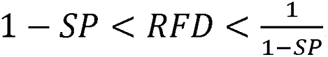. If this inequality is reversed, i.e. 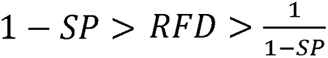, the two species display positive frequency dependence (Ke and Letten 2018; Zou and Rudolf 2023), i.e., the species with a higher initial abundance wins.

We conducted all analyses in R 4.2.1 (R Core Team 2024) and fitted the Bayesian model in Stan using the package rstan (Stan Development Team 2024). We fitted all models with a total of 100,000 iterations, using the first 50,000 as burn-in. All models converged well (all rhat values < 1.001). Data and code are available at https://github.com/hengxingzou/Zou2025bioRXiv/ and will be archived to a permanent repository upon acceptance.

## Results

### Final Population Size

In general, the earlier one species arrives, the larger its final adult population, indicating strong priority effects (Figure 2; Figure S3-S5). Without the competing species, *T. confusum* reached higher densities than *T. castaneum*, suggesting overall higher fecundity (Table S1, S3), but *T. castaneum* appeared competitively dominant because it could completely exclude *T. confusum* when arriving early (Figure S3-S5). For both species, the best-fitting linear mixed effects model included the number of generations, relative arrival times, and the ratio of initial egg densities, indicating that adult populations are strongly affected by all three factors (Table S2). The models did not detect significant random effect across replicates of each treatment.

**Figure 2.**
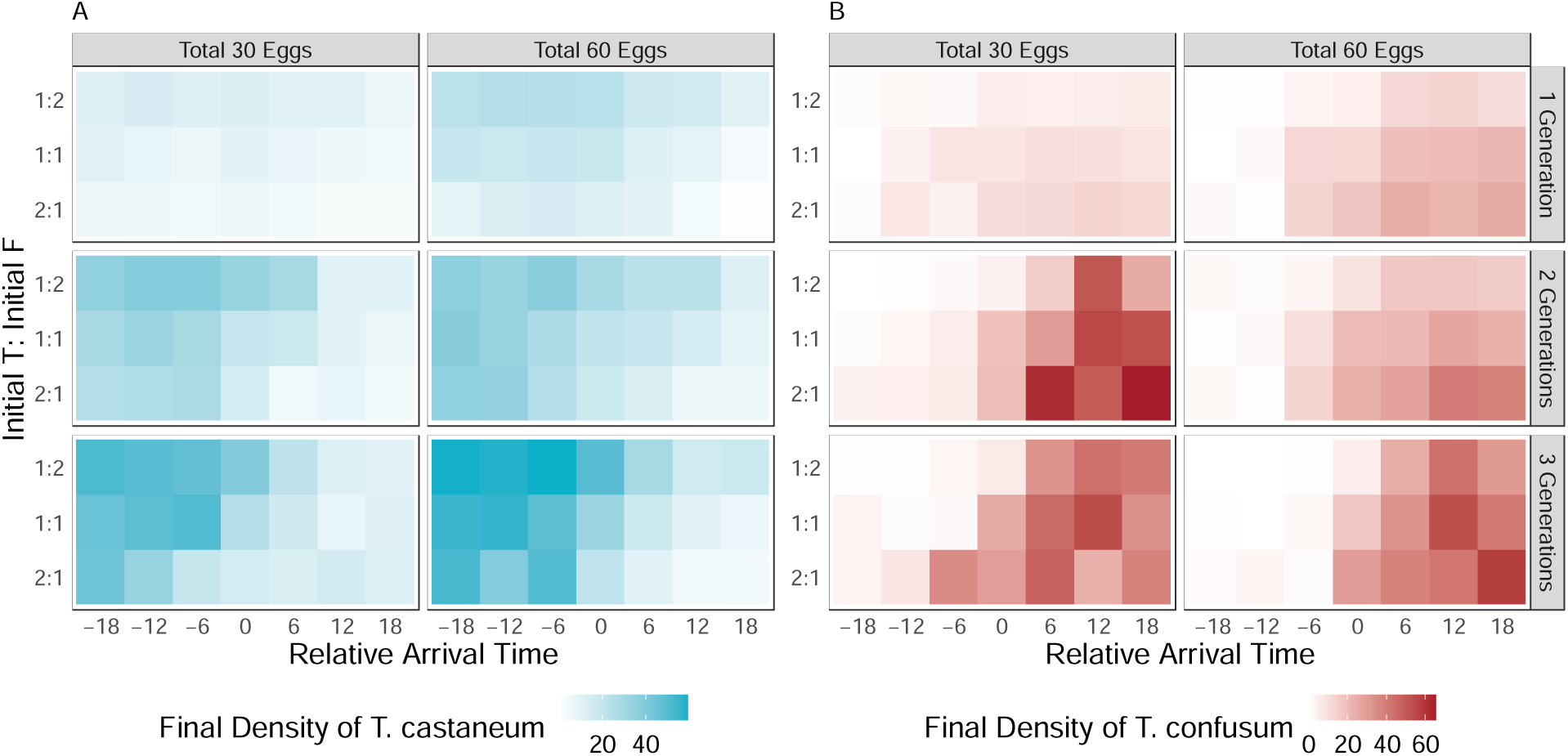
Final adult populations in vials with two species (A. *T. castaneum*; B. *T. confusum*), by total densities, the ratio of initial egg densities, the number of generations and relative arrival times. Adult population is averaged across the three replicates. Darker shade indicates higher adult population.

Priority effects were reflected by the end populations: with one generation per season, the end population of both species decreased when it arrived later (*T. castaneum*: estimated slope = - 0.79, SE = 0.038; *T. confusum*: estimated slope = 0.99, SE = 0.049; both *p* < 0.01; slopes differ in sign because negative relative arrival times indicate early arrival of *T. castaneum*, and positive relative arrival times indicate early arrival of *T. confusum*). More generations in a season led to higher total populations of both species (*T. castaneum*: estimated slope = 9.77, SE = 0.56; *T. confusum*: estimated slope = 6.28, SE = 0.72; both *p* < 0.01). For both species, an initial numeric advantage led to up to 10-fold increases of final population: increasing the initial relative abundance from 1:2 to 1:1 and 2:1 increased the final population of *T. castaneum* by 4.94 (SE = 1.12, *p* < 0.01) and by 10.57 (SE = 1.12, *p* < 0.01) and for *T. confusum* by 5.97 (SE = 1.45, *p* < 0.01), and by 9.86 (SE = 1.45, *p* < 0.01), respectively (see Table S2 for full list of statistics).

### Competition Coefficients

Fitted interspecific per-capita competition coefficients further illustrated the presence of priority effects. The earlier one species arrives, the stronger its per-capita competitive effect towards the late arriver (Figure 3). The magnitude of the two competition coefficients flipped at a positive relative arrival time (around 6 days), i.e., *T. confusum* needed to arrive earlier to achieve equal interspecific competition between *T. castaneum*. This suggests that *T. castaneum* was competitively dominant in general. This priority effect was consistent with the differences in predation of eggs across species we observed in our predation trials: the larvae of *T. castaneum* had a higher predation rate on the eggs of *T. confusum*, with larger larvae consuming more eggs (Figure S7). However, the number of generations did not have a consistent effect on the strength of priority effects: although the two competition coefficients were more similar when *T. confusum* arrived earlier with two generations per season, this similarity was absent with three generations per season.

**Figure 3.**
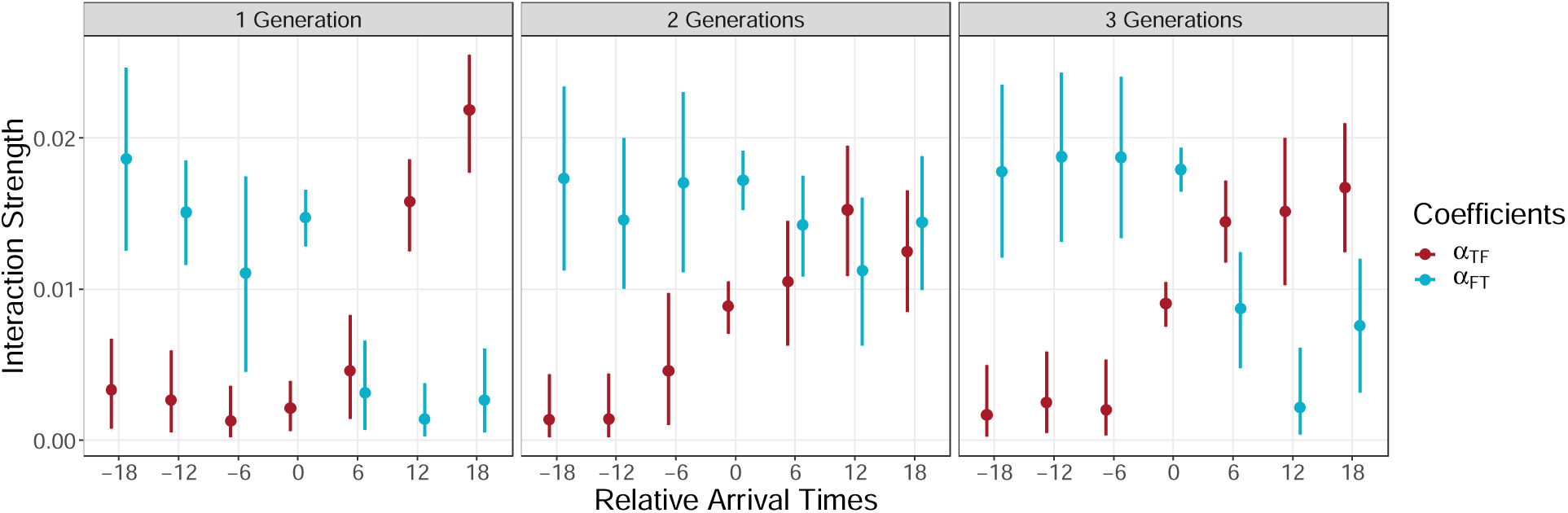
Fitted interspecific competition coefficients in a discrete-time Lotka-Volterra model. α*_TF_* indicates the effect from *T. confusum* to *T. castaneum*, and vice versa. A negative relative arrival time indicates the early arrival of *T. castaneum*, while a 0 indicates simultaneous arrival. Points show the median (50%), and line ranges show 10% and 90% of the posterior distribution.

### Niche and Fitness Differences

Calculated stabilization potential and relative fitness differences showed much larger variations across different arrival times with one vs. two or three generations per season (Figure 4). With one generation per season, mean stabilization potentials were always negative and, combined with strong relative fitness differences, led to a strong tendency towards exclusion when either species arrives early. At simultaneous arrival, the two species tended to display positive frequency dependence. With more generations per season, both niche and fitness differences between the species were much smaller but still generally predicted exclusion when either species arrived early. At simultaneous arrival, *T. castaneum* tended to exclude *T. confusum*, matching our expectation that the former was competitively superior. With two generations per season, when *T. confusum* arrived early by 12 and 18 days, the overall niche and fitness differences predicted frequency-dependent priority effects, a result different from both one and three generations per season. This likely arose from the particularly high population of *T. confusum* when it arrived early and has a numeric advantage (Figure S3-S5), which affected the fitted competition coefficients and, subsequently, stabilization potentials and fitness differences. With three generations per season, the early arrival of either species led to more positive niche difference values, suggesting a stabilizing effect of arriving at different times. However, the two species were still predicted to exclude each other due to the high fitness differences.

**Figure 4.**
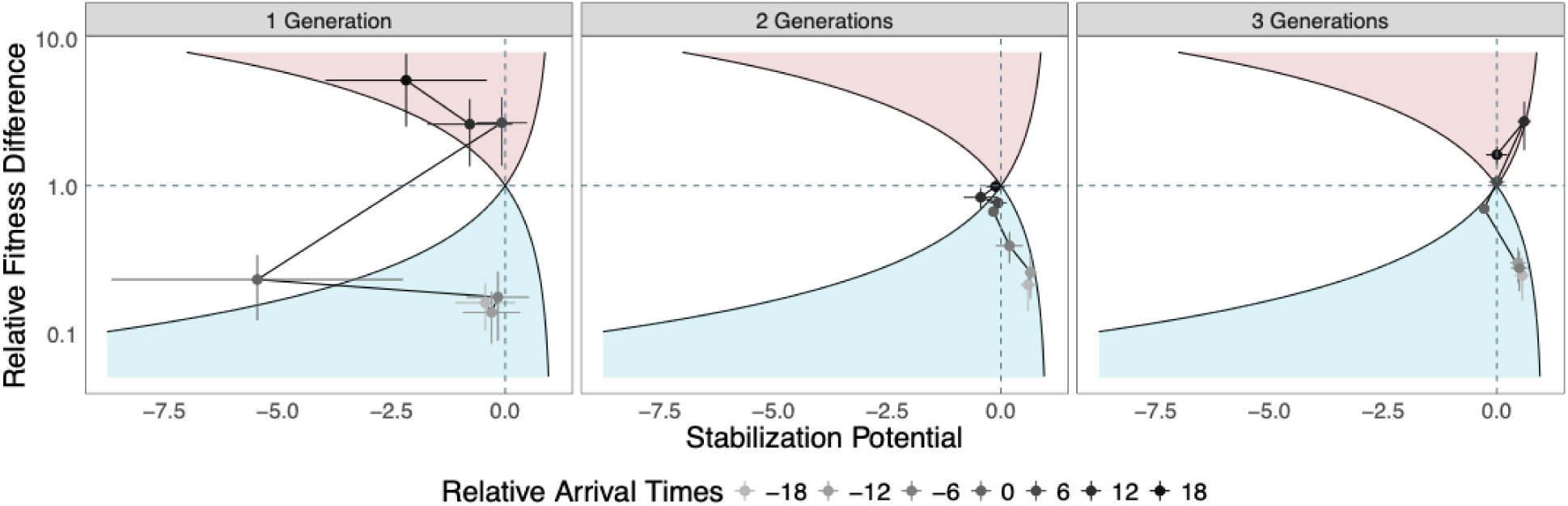
Stabilization potentials and relative fitness differences calculated from 1000 randomly sampled posterior estimates of intra- and interspecific competition coefficients. Points and error bars indicate the means of calculated ND and RFD values plus and minus one standard deviation. Blue shaded region below the solid curves indicates *T. castaneum* wins, red shaded region above the solid curves indicates *T. confusum* wins. The region to the left of both curves indicates frequency-dependent priority effects. The region to the right indicates coexistence. Points show values calculated by the median, and line ranges show values calculated by 10% and 90% of the posterior distribution of the competition coefficients.

## Discussion

As we strive to predict consequences of phenological shifts across systems, understanding the role of life history in modulating these effects is fundamental. We explored how one important life history trait, the number of generations per growing season, shapes the competitive dynamics between two flour beetles as they arrive at different times. We found strong changes in interspecific competition with changes in relative arrival time, indicating trait-dependent priority effects. Furthermore, more generations lead to increasingly positive stabilization potentials and smaller relative fitness differences between the two species, indicating a decrease in such trait-dependent priority effects. These results show the nuanced relationship between the consequence of phenological difference and the number of generations.

### Separating Different Mechanisms of Priority Effects

Increasing experimental evidence suggests that priority effects can arise from changes in species interactions over their arrival times, often due to changes in traits that affect these biotic interactions (Shorrocks and Bingley 1994; Rudolf 2018; Blackford et al. 2020; Fragata et al. 2022). We observed changes in competition coefficients over the relative arrival times between the two flour beetles, indicating strong trait-dependent priority effects (Zou and Rudolf 2023). All stages of flour beetles compete readily for resources, and the later stages (larger larvae and adults) should consume resources at higher rates than smaller larvae (Maino and Kearney 2015). Because the earlier arriving species had more time to increase in relative size/stage, it gained a competitive advantage, leading to trait-dependent priority effects (Zou et al. 2023). In addition to resource competition, larvae of the early arriver may consume eggs of the late arriver at the time of the egg addition, further enhancing priority effects. Indeed, we found that larger larvae of *T. castaneum* consumed more eggs of the late-arriving *T. confusum*, but not vice versa, leading to an even stronger early arriver advantage for *T. castaneum* through intraguild predation. When combined with other effects such as size-mediated resource competition, *T. castaneum* could drive *T. confusum* extinct in just one generation. Together, these mechanisms strengthened trait-dependent priority effects between the two species, as indicated by both fitted competition coefficients and calculated niche and fitness differences across relative arrival times.

In addition to these mechanisms, priority effects can also arise from a higher abundance of the early arriving species, which can be further maintained by positive frequency dependence (i.e., frequency-dependent priority effects; Ke and Letten 2018; Toju et al. 2018; Zou and Rudolf 2023). Indeed, previous experiments on *Tribolium* beetles found that the species with higher initial abundance usually wins (Park 1957). In our experiments, the initial frequency of the two flour beetles affected the final adult population, indicating the presence of frequency dependence. Stabilization potentials and relative fitness differences further suggested that frequency-dependent priority effects were likely to occur when the two species arrived simultaneously with one generation per season or when *T. confusum* arrived very early with two generations per season. These results indicate the prevalence of trait-dependent priority effects (Fragata et al. 2022) and that the two different mechanisms (trait-dependent and frequency-dependent) of priority effects can concurrently shape community dynamics.

### Nuanced Effects of the Number of Generations on Phenological Differences

Previous models predicted that more overlapping generations in a season should reduce the difference in stage distributions of the two species and thus lower the strength of trait-dependent priority effects (Zou et al. 2023). Yet, our experiment found a more nuanced response. To understand the discrepancies, we need to identify the possible biological mechanisms that caused the signs and strengths of phenomenological stabilization potential and relative fitness differences. A higher stabilization potential indicates that species limit themselves more than each other (e.g., via resource partitioning), which should promote stable coexistence. A negative stabilization potential indicates that species limit each other more than themselves, which promotes alternative stable states of competitive exclusion (see Ke and Letten 2018 for a sematic discussion on the terminology). On the other hand, smaller fitness difference “equalizes” the competitive ability between two species, promoting coexistence (Chesson 2000, 2018). In our experiment, the strongly negative stabilization potential with one generation per season indicated the high ecological similarity between *T. castaneum* and *T. confusum*. However, stabilization potentials became less negative and even switched to positive values with longer seasons, suggesting that arriving at different times promoted the differentiation between the two species. One possible source of such differentiation is that a large initial difference in arrival times may exempt the larvae of late-arriving species from some resource competition when the early-arriving species pupated and did not forage. However, this difference in size/stage distribution should decline with more overlapping generations, yet we observed increasing stabilization potential instead, suggesting other unaccounted mechanisms.

On the other hand, some biological characteristics led to fitness differences between the two species. *T. castaneum* develops faster (Park 1948, 1954) and, in our experiment, consumed more eggs of *T. confusum*. *T. confusum* always reached higher population in single-species vials due to its higher fecundity in our experiment and higher tolerance to unfavorable environmental conditions (high population density, low availability of resources, or flour previously “conditioned” by other beetles; Riddle 1976; Riddle et al. 1986). With only one generation, the fast development and high egg predation rate made *T. castaneum* competitively dominant, sometimes leading to the extinction of the *T. confusum*. This explains why we observed strong fitness differences between the two species with only one generation. However, if enough *T. confusum* survived to adulthood after the first generation (more likely when *T. confusum* is initially more abundant), their higher fecundity could buffer against the high predation rate of their eggs by *T. castaneum*, leading to apparent coexistence in the experimental vials. Therefore, more generations per season led to smaller fitness differences and weaker competitive asymmetry. This buffering might not increase in strength with more generations because it mostly depends on the number of surviving *T. confusum* adults after the first generation. This could explain why patterns in niche and fitness differences only changed qualitatively from one to two generations per season, but not from two to three generations per season, contrary to theoretical predictions that more generations per season should lead to even weaker trait-dependent priority effects (Zou et al. 2023).

### Implications and Future Directions

Despite extensive experimental and theoretical explorations, few studies consider the long-term consequences of priority effects in the context of repeated disturbance followed by re-assembly of communities (Zou and Rudolf 2023). This periodic reassembly is widespread in seasonal ecosystems and may lead to compositional cycles or alternative transient states, and the long-term dynamics depend on the strengths of species interactions (Song et al. 2021; Spaak and Schreiber 2023; Song 2025). Therefore, exploring the effects of key processes of seasonal communities, such as dormancy (Wisnoski et al. 2019), stochasticity (Stump and Vasseur 2023), and reproduction regimes on species interactions is important to predict long-term dynamics of ecological communities. Our experiment focuses on the reproduction regime within a growing season and provides a first empirical test of recent theoretical exploration of the interaction between the number of generations per season and the strengths of trait-dependent priority effects (Zou et al. 2023). Instead of the increasing dominance of frequency-dependent priority effects predicted by the theoretical model, we found sustained but weakened trait-dependent priority effects in our experiments. This deviation might arise from the relatively shorter duration of our experiment, but more importantly, it shows that even in a controlled system, empirical systems may contain biotic processes that are unaccounted for in theoretical models. Given the prevalence and importance of seasonal communities in nature, our results further highlight the importance of testing theoretical predictions of periodic reassembly by experiments.

Life history traits, such as the number of reproductive events per growing season, can be highly plastic in response to environmental changes (Altermatt 2010; Brans and De Meester 2018; Keller and Shea 2020). Studying the community-level consequences of these shifts is therefore fundamental to understanding nature’s responses to climate change. Our experiment indicates that the consequences of phenological shifts may change in systems with different numbers of generations in a growing season. However, our experiment only simulated the shift in the reproduction regime by manually imposing the length of a growing season. Further experiments that can induce such changes in life history by directly manipulating the environment are needed to better characterize the interaction between life history change and priority effects. Together, our results can assist the development of a unified theory of priority effects across different communities and the evaluation of priority effects under the synergistic shifts of phenology and life history under climate change.

### Author Contributions

H.-X.Z. conceived the idea and developed the methods with assistance from V.H.W.R. H.-X.Z. conducted the experiment, performed analyses with assistance from V.H.W.R., and wrote the first draft. Both authors contributed to revisions.

## Acknowledgement

We thank Ann T. Tate and Jeff P. Demuth for their help and suggestions on the beetles’ maintenance protocol. We thank Tom E.X. Miller, Ali Campbell, Jacob K. Moutouama, and Daniel Kowal for feedback on the manuscript and help with statistical methods. Funding was provided by NSF DEB-1655626.

