## Supplementary Material for "Consequences of phenological shifts are determined by the number of generations per season"

**Bayesian Model Fitting**

We fit the final count of the adults of species *i* with number of generations *G* as:

$C\left( t \right)_{i,G}\sim Normal\left( N_{i,G},\sigma\right)$ (S.1).

We used a normal distribution because within each treatment of the number of generations, fitting the model with a Poisson or negative binomial distribution led to convergence issues (>1000 divergent transitions after the warmup). We then fitted two alternative models, Beverton-Holt (Eqn. S2) and Ricker (Eqn. S3), to one dataset from our experiments (single-species vials of *T. castaneum*, one generation, *n*=6):

$\frac{N\left( t \right)_{i,G}}{N\left( 0 \right)_{i,G}}=\frac{\lambda_{i,G}}{1+\alpha_{ii,G}N\left( 0 \right)_{i,G}}$ (S.2);

$\frac{N\left( t \right)_{i,G}}{N\left( 0 \right)_{i,G}}=\exp\left( \lambda_{i,G}\left( 1-\alpha_{ii,G}N\left( 0 \right)_{i,G} \right) \right)$ (S.3).

We then calculated the average log likelihood of the posterior. Both Beverton-Holt and Ricker models had convergence issues and lower log likelihood (-4.012 and -4.502) than a discrete Lotka-Volterra model (-2.926). We therefore chose the Lotka-Volterra model (Eqn. 1 in the main text) to fit all the datasets.

To further evaluate the goodness of model fitting, we compared the fitted growth rate, $\hat{\frac{N\left( t \right)_{i,G}}{N\left( 0 \right)_{i,G}}}=\hat{\lambda_{i,G}}\left( 1-\hat{\alpha_{ii,G}}N\left( 0 \right)_{i,G} \right)$, where parameters with the hat are fitted from the model, to the actual growth rate calculated from experimental data. Fitted growth rates generally match well with the actual data (Figure S1, S2), except when one species is extremely favored/unfavored. For instance, when *T. castaneum* arrives early its actual growth rate can greatly exceed the fitted growth rate, at which the actual growth rate of *T. confusum* is lower than fitted. This is because the advantage of *T. castaneum* was so large that population of *T. confusum* was close to 0.

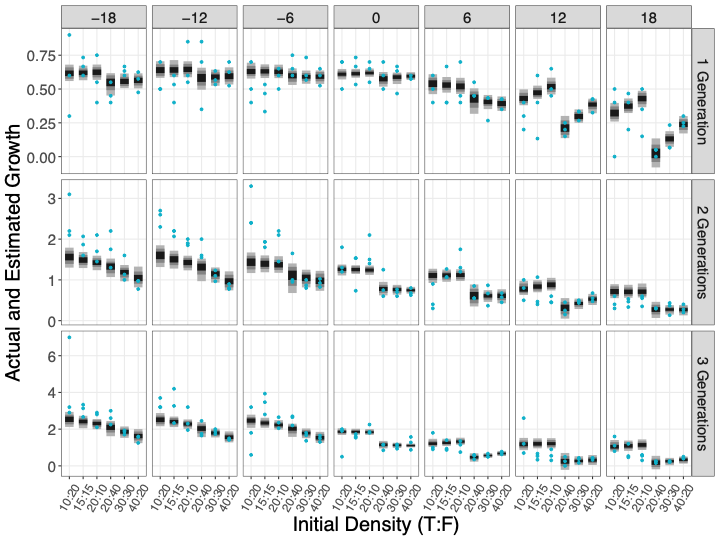
**Figure S1.** Actual and fitted growth rates ($N\left( t \right)/N\left( 0 \right)$) of *T. castaneum*. Rows represent number of generations and columns represent relative arrival times. Shaded intervals show growth rates calculated from 1000 randomly sampled sets of parameters, and darker to lighter shades show the 50%, 80%, and 95% rates around the median. Blue points show actual growth rates in the experiments. Initial egg densities (*T. castaneum* to *T. confusum*) are shown on the x axis.

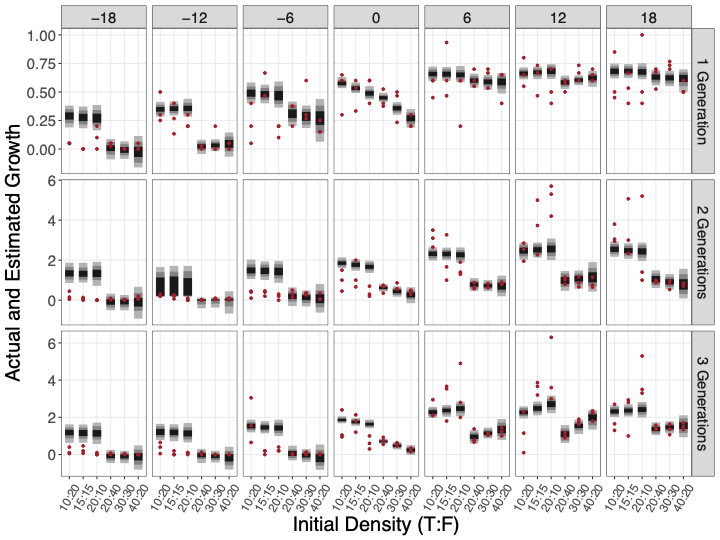

**Figure S2.** Actual and fitted growth rates ($N\left( t \right)/N\left( 0 \right)$) of *T. confusum*. Rows represent number of generations and columns represent relative arrival times. Shaded intervals show growth rates calculated from 1000 randomly sampled sets of parameters, and darker to lighter shades show the 50%, 80%, and 95% rates around the median. Red points show actual growth rates in the experiments. Initial egg densities (*T. castaneum* to *T. confusum*) are shown on the x axis.

**Final Adult Population**

| **Species** | **Number of Generations** | **Initial Population** | **Average Final Population** | **Standard Error (n=3)** |
| --- | --- | --- | --- | --- |
| *T. castaneum* | 1 | 30 | 15.67 | 0.88 |
|  |  | 60 | 38.33 | 1.20 |
|  | 2 | 30 | 34.67 | 0.88 |
|  |  | 60 | 43.33 | 0.88 |
|  | 3 | 30 | 56.00 | 3.21 |
|  |  | 60 | 62.00 | 3.21 |
| *T. confusum* | 1 | 30 | 20.00 | 2.00 |
|  |  | 60 | 38.00 | 1.15 |
|  | 2 | 30 | 71.00 | 6.11 |
|  |  | 60 | 57.33 | 4.05 |
|  | 3 | 30 | 68.67 | 5.55 |
|  |  | 60 | 69.00 | 1.73 |

**Table S1.** Final densities of single-species vials.

| **Species** | **Variable** | **Estimate** | **Standard Error** | **Degree of Freedom** | **t Value** | ***p*** |
| --- | --- | --- | --- | --- | --- | --- |
| *T. castaneum* | (Intercept) | -3.55 | 1.37 | 373 | -2.58 | 0.010 |
|  | Number of Generations | 9.77 | 0.56 | 373 | 17.43 | **< 2e-16** |
|  | Relative Arrival Times | -0.78 | 0.038 | 373 | -20.42 | **< 2e-16** |
|  | Ratio 1 (1:1) | 4.94 | 1.12 | 373 | 4.41 | **1.35e-5** |
|  | Ratio 2 (2:1) | 10.57 | 1.12 | 373 | 9.43 | **<2e-16** |
|  | **Random Effects:** | | | | | |
|  | Group | Variance | Standard Deviation |  |  |  |
|  | Replication | 0.00 | 0.00 |  |  |  |
|  | Residual | 79.16 | 8.897 |  |  |  |
|  | N = 378, groups: Replication, 3 | | | | | |
|  | ∆AIC from the second best-fit model: -146.30 | | | | | |
| *T. confusum* | (Intercept) | 8.28 | 1.78 | 373 | 4.66 | 4.40e-6 |
|  | Number of Generations | 6.28 | 0.73 | 373 | 8.66 | **< 2e-16** |
|  | Relative Arrival Times | 0.99 | 0.049 | 373 | 20.02 | **< 2e-16** |
|  | Ratio 1 (1:1) | -3.89 | 1.45 | 373 | -2.68 | **0.0077** |
|  | Ratio 2 (2:1) | -9.86 | 1.45 | 373 | -6.80 | **4.25e-11** |
|  | **Random Effects:** | | | | | |
|  | Group | Variance | Standard Deviation |  |  |  |
|  | Replication | 0.00 | 0.00 |  |  |  |
|  | Residual | 132.5 | 11.51 |  |  |  |
|  | N = 378, groups: Replication, 3 | | | | | |
|  | ∆AIC from the second best-fit model: -26.50 | | | | | |

**Table S2.** Summary statistics of the linear mixed effects model for the final adult population of each species. Ratio represents the ratio of initial egg densities (1:1, 1:2, 2:1) and is always in the order of *T. castaneum* : *T. confusum*.

| **Relative Arrival Time** | **Number of Generation** | **Initial Egg Ratio (*T*:*F*)** | **Total Number of Eggs** | **Average Final Population, *T. castaneum*** | **Standard Error, *T. castaneum* (n=3)** | **Average Final Population, *T. confusum*** | **Standard Error, *T. confusum* (n=3)** |
| --- | --- | --- | --- | --- | --- | --- | --- |
| 0 | 1 | 0.5 | 30 | 6.33333333 | 0.66666667 | 10.3333333 | 2.18581284 |
| 0 | 1 | 1 | 30 | 9.66666667 | 0.8819171 | 7.33333333 | 1.20185043 |
| 0 | 1 | 2 | 30 | 12.6666667 | 0.8819171 | 5.33333333 | 0.66666667 |
| 12 | 1 | 0.5 | 30 | 3 | 0.57735027 | 13.3333333 | 1.45296631 |
| 12 | 1 | 1 | 30 | 5.66666667 | 2.02758751 | 9.33333333 | 1.20185043 |
| 12 | 1 | 2 | 30 | 10.6666667 | 1.20185043 | 5.33333333 | 0.8819171 |
| 18 | 1 | 0.5 | 30 | 3 | 1.52752523 | 12 | 2.51661148 |
| 18 | 1 | 1 | 30 | 5.33333333 | 1.20185043 | 8 | 1.15470054 |
| 18 | 1 | 2 | 30 | 6.66666667 | 2.02758751 | 6.33333333 | 1.85592145 |
| 6 | 1 | 0.5 | 30 | 5 | 0.57735027 | 11 | 1 |
| 6 | 1 | 1 | 30 | 7.33333333 | 1.33333333 | 10 | 2.081666 |
| 6 | 1 | 2 | 30 | 10.3333333 | 1.85592145 | 4.66666667 | 1.33333333 |
| -12 | 1 | 0.5 | 30 | 5.66666667 | 0.66666667 | 7 | 1.52752523 |
| -12 | 1 | 1 | 30 | 7.66666667 | 0.8819171 | 4 | 1.15470054 |
| -12 | 1 | 2 | 30 | 13.3333333 | 1.85592145 | 2.33333333 | 0.33333333 |
| -18 | 1 | 0.5 | 30 | 6 | 1.73205081 | 1 | 0 |
| -18 | 1 | 1 | 30 | 10 | 0.57735027 | 0 | 0 |
| -18 | 1 | 2 | 30 | 11.3333333 | 2.02758751 | 1 | 0.57735027 |
| -6 | 1 | 0.5 | 30 | 5.33333333 | 0.8819171 | 4.33333333 | 2.02758751 |
| -6 | 1 | 1 | 30 | 6.66666667 | 0.8819171 | 8 | 1 |
| -6 | 1 | 2 | 30 | 11.6666667 | 0.8819171 | 1.66666667 | 0.33333333 |
| 0 | 1 | 0.5 | 60 | 12 | 1.15470054 | 17.3333333 | 1.85592145 |
| 0 | 1 | 1 | 60 | 17.6666667 | 1.85592145 | 12 | 2.51661148 |
| 0 | 1 | 2 | 60 | 23.3333333 | 0.33333333 | 4.66666667 | 0.66666667 |
| 12 | 1 | 0.5 | 60 | 4 | 0.57735027 | 22.3333333 | 1.20185043 |
| 12 | 1 | 1 | 60 | 9.33333333 | 0.66666667 | 20 | 1.15470054 |
| 12 | 1 | 2 | 60 | 15.6666667 | 1.33333333 | 13 | 0.57735027 |
| 18 | 1 | 0.5 | 60 | 0.66666667 | 0.33333333 | 25.3333333 | 2.18581284 |
| 18 | 1 | 1 | 60 | 3.66666667 | 1.66666667 | 22 | 0.57735027 |
| 18 | 1 | 2 | 60 | 10.3333333 | 0.8819171 | 10.6666667 | 0.66666667 |
| 6 | 1 | 0.5 | 60 | 9.66666667 | 0.66666667 | 24.6666667 | 1.76383421 |
| 6 | 1 | 1 | 60 | 11.3333333 | 1.66666667 | 19 | 1.52752523 |
| 6 | 1 | 2 | 60 | 16 | 1 | 11.3333333 | 1.66666667 |
| -12 | 1 | 0.5 | 60 | 12.6666667 | 2.96273147 | 0.66666667 | 0.33333333 |
| -12 | 1 | 1 | 60 | 17.3333333 | 0.8819171 | 2 | 2 |
| -12 | 1 | 2 | 60 | 24.6666667 | 2.02758751 | 0.66666667 | 0.33333333 |
| -18 | 1 | 0.5 | 60 | 9.33333333 | 0.8819171 | 1.66666667 | 0.33333333 |
| -18 | 1 | 1 | 60 | 19 | 0.57735027 | 0 | 0 |
| -18 | 1 | 2 | 60 | 22.3333333 | 1.76383421 | 0.33333333 | 0.33333333 |
| -6 | 1 | 0.5 | 60 | 13.3333333 | 0.8819171 | 12.6666667 | 2.33333333 |
| -6 | 1 | 1 | 60 | 19 | 1.52752523 | 11.6666667 | 3.17979734 |
| -6 | 1 | 2 | 60 | 23.6666667 | 1.45296631 | 3.66666667 | 0.66666667 |
| 0 | 2 | 0.5 | 30 | 14.3333333 | 1.85592145 | 19.6666667 | 6.06446847 |
| 0 | 2 | 1 | 30 | 19 | 4 | 18.3333333 | 6.00925213 |
| 0 | 2 | 2 | 30 | 33.3333333 | 4.37162568 | 4 | 1.52752523 |
| 12 | 2 | 0.5 | 30 | 7.66666667 | 1.45296631 | 49 | 5.29150262 |
| 12 | 2 | 1 | 30 | 9.66666667 | 3.17979734 | 55 | 11.8462371 |
| 12 | 2 | 2 | 30 | 10 | 1 | 50.6666667 | 4.48454135 |
| 18 | 2 | 0.5 | 30 | 4.33333333 | 0.8819171 | 65.3333333 | 5.36449231 |
| 18 | 2 | 1 | 30 | 6.66666667 | 0.8819171 | 52 | 12.3423391 |
| 18 | 2 | 2 | 30 | 10 | 1.52752523 | 25.3333333 | 13.3832399 |
| 6 | 2 | 0.5 | 30 | 5.33333333 | 1.85592145 | 61.6666667 | 4.91030662 |
| 6 | 2 | 1 | 30 | 17 | 1 | 29.6666667 | 10.0884973 |
| 6 | 2 | 2 | 30 | 28.3333333 | 3.38296386 | 15.3333333 | 1.85592145 |
| -12 | 2 | 0.5 | 30 | 25.3333333 | 1.20185043 | 4.66666667 | 0.66666667 |
| -12 | 2 | 1 | 30 | 32.3333333 | 0.66666667 | 2.66666667 | 0.8819171 |
| -12 | 2 | 2 | 30 | 38.3333333 | 0.8819171 | 0.33333333 | 0.33333333 |
| -18 | 2 | 0.5 | 30 | 24.6666667 | 3.17979734 | 4.33333333 | 2.40370085 |
| -18 | 2 | 1 | 30 | 27.6666667 | 2.02758751 | 1 | 0.57735027 |
| -18 | 2 | 2 | 30 | 35 | 3.7859389 | 0 | 0 |
| -6 | 2 | 0.5 | 30 | 27 | 3 | 6.33333333 | 2.18581284 |
| -6 | 2 | 1 | 30 | 28.3333333 | 0.66666667 | 5.33333333 | 1.20185043 |
| -6 | 2 | 2 | 30 | 38.3333333 | 4.70224533 | 1.66666667 | 0.8819171 |
| 0 | 2 | 0.5 | 60 | 17.3333333 | 3.92994204 | 23.6666667 | 4.91030662 |
| 0 | 2 | 1 | 60 | 20.6666667 | 1.45296631 | 20 | 3 |
| 0 | 2 | 2 | 60 | 28 | 2.081666 | 8.33333333 | 1.20185043 |
| 12 | 2 | 0.5 | 60 | 5.66666667 | 1.76383421 | 39.6666667 | 4.09606858 |
| 12 | 2 | 1 | 60 | 14 | 0.57735027 | 27 | 4.35889894 |
| 12 | 2 | 2 | 60 | 22.6666667 | 2.33333333 | 17 | 3.05505046 |
| 18 | 2 | 0.5 | 60 | 6 | 0 | 37.6666667 | 1.33333333 |
| 18 | 2 | 1 | 60 | 9.33333333 | 2.72845092 | 23.3333333 | 3.84418753 |
| 18 | 2 | 2 | 60 | 11.6666667 | 2.33333333 | 15.6666667 | 0.8819171 |
| 6 | 2 | 0.5 | 60 | 13 | 2 | 28 | 3.60555128 |
| 6 | 2 | 1 | 60 | 19 | 4.35889894 | 21 | 2 |
| 6 | 2 | 2 | 60 | 22.6666667 | 2.60341656 | 16.6666667 | 0.66666667 |
| -12 | 2 | 0.5 | 60 | 34.3333333 | 2.84800125 | 0.66666667 | 0.33333333 |
| -12 | 2 | 1 | 60 | 33 | 2.30940108 | 2 | 0.57735027 |
| -12 | 2 | 2 | 60 | 33 | 1.52752523 | 1 | 0.57735027 |
| -18 | 2 | 0.5 | 60 | 35 | 5.19615242 | 2.33333333 | 0.66666667 |
| -18 | 2 | 1 | 60 | 37.3333333 | 5.45690185 | 0.33333333 | 0.33333333 |
| -18 | 2 | 2 | 60 | 36.3333333 | 2.66666667 | 2.33333333 | 0.8819171 |
| -6 | 2 | 0.5 | 60 | 24 | 4.50924975 | 12.6666667 | 4.05517502 |
| -6 | 2 | 1 | 60 | 26.6666667 | 1.76383421 | 9.66666667 | 0.8819171 |
| -6 | 2 | 2 | 60 | 37 | 2.30940108 | 3 | 2 |
| 0 | 3 | 0.5 | 30 | 14.6666667 | 4.84194635 | 29.3333333 | 9.35117343 |
| 0 | 3 | 1 | 30 | 24.3333333 | 0.8819171 | 25.6666667 | 4.09606858 |
| 0 | 3 | 2 | 30 | 39.3333333 | 2.84800125 | 6.33333333 | 2.02758751 |
| 12 | 3 | 0.5 | 30 | 15 | 5.6862407 | 23.6666667 | 12.706079 |
| 12 | 3 | 1 | 30 | 7.66666667 | 1.45296631 | 53.6666667 | 2.96273147 |
| 12 | 3 | 2 | 30 | 11.6666667 | 3.48010217 | 43 | 10.1488916 |
| 18 | 3 | 0.5 | 30 | 11.6666667 | 2.33333333 | 37.6666667 | 8.41295298 |
| 18 | 3 | 1 | 30 | 11 | 3.51188458 | 33.6666667 | 9.35117343 |
| 18 | 3 | 2 | 30 | 9.66666667 | 1.85592145 | 40.3333333 | 6.35959468 |
| 6 | 3 | 0.5 | 30 | 12.3333333 | 2.84800125 | 47.6666667 | 5.66666667 |
| 6 | 3 | 1 | 30 | 16 | 2.51661148 | 45 | 9.01849951 |
| 6 | 3 | 2 | 30 | 21.6666667 | 3.52766841 | 32.3333333 | 8.64741451 |
| -12 | 3 | 0.5 | 30 | 33.6666667 | 1.66666667 | 7.33333333 | 3.48010217 |
| -12 | 3 | 1 | 30 | 49 | 8.08290377 | 1 | 1 |
| -12 | 3 | 2 | 30 | 49.6666667 | 7.44610263 | 0.66666667 | 0.33333333 |
| -18 | 3 | 0.5 | 30 | 43.6666667 | 13.1951169 | 3.66666667 | 2.18581284 |
| -18 | 3 | 1 | 30 | 45.6666667 | 2.96273147 | 3.66666667 | 1.76383421 |
| -18 | 3 | 2 | 30 | 52 | 5.03322296 | 0.33333333 | 0.33333333 |
| -6 | 3 | 0.5 | 30 | 18.6666667 | 7.51295178 | 35 | 14 |
| -6 | 3 | 1 | 30 | 51 | 4.93288286 | 2 | 1 |
| -6 | 3 | 2 | 30 | 47 | 3.46410162 | 2.66666667 | 0.66666667 |
| 0 | 3 | 0.5 | 60 | 21 | 2 | 29.6666667 | 4.33333333 |
| 0 | 3 | 1 | 60 | 31.6666667 | 2.02758751 | 17 | 1.52752523 |
| 0 | 3 | 2 | 60 | 49.6666667 | 8.11035004 | 5 | 1.15470054 |
| 12 | 3 | 0.5 | 60 | 5 | 2.88675135 | 41 | 3.21455025 |
| 12 | 3 | 1 | 60 | 9.66666667 | 2.40370085 | 53 | 2.30940108 |
| 12 | 3 | 2 | 60 | 15.6666667 | 1.33333333 | 42.6666667 | 3.38296386 |
| 18 | 3 | 0.5 | 60 | 4.33333333 | 0.66666667 | 57.6666667 | 2.72845092 |
| 18 | 3 | 1 | 60 | 7 | 0.57735027 | 40.6666667 | 4.91030662 |
| 18 | 3 | 2 | 60 | 17.3333333 | 1.33333333 | 30.3333333 | 2.02758751 |
| 6 | 3 | 0.5 | 60 | 10 | 1.52752523 | 37.6666667 | 8.74325137 |
| 6 | 3 | 1 | 60 | 16.6666667 | 1.66666667 | 32.6666667 | 0.8819171 |
| 6 | 3 | 2 | 60 | 28.3333333 | 1.76383421 | 24.6666667 | 2.60341656 |
| -12 | 3 | 0.5 | 60 | 39 | 5.03322296 | 3 | 1.52752523 |
| -12 | 3 | 1 | 60 | 56 | 2.30940108 | 0 | 0 |
| -12 | 3 | 2 | 60 | 58 | 1 | 0 | 0 |
| -18 | 3 | 0.5 | 60 | 52.3333333 | 4.33333333 | 1.33333333 | 0.8819171 |
| -18 | 3 | 1 | 60 | 55 | 2.081666 | 0.66666667 | 0.33333333 |
| -18 | 3 | 2 | 60 | 59 | 4.50924975 | 0 | 0 |
| -6 | 3 | 0.5 | 60 | 50.3333333 | 3.17979734 | 1 | 0.57735027 |
| -6 | 3 | 1 | 60 | 48.3333333 | 3.84418753 | 2.33333333 | 1.20185043 |
| -6 | 3 | 2 | 60 | 59.3333333 | 4.05517502 | 0.33333333 | 0.33333333 |

**Table S3.** Final densities of two-species vials. Negative relative arrival times means that *T. castaneum* arrives early.

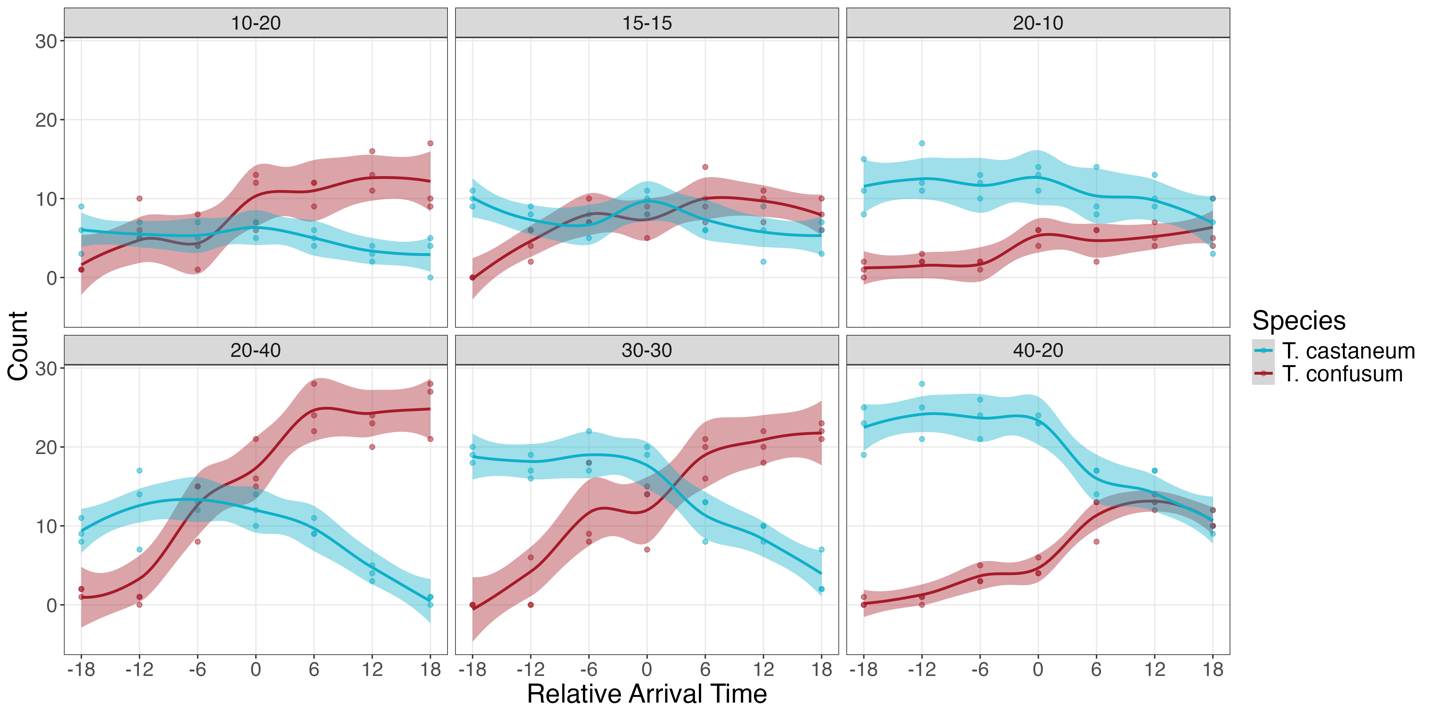

**Figure S3.** Final adult populations of vials with one generation. Numbers on top of each panel shows the initial population in the order of *T. castaneum* – *T. confusum*. Negative relative arrival times represents the early arrival of *T. castaneum*.

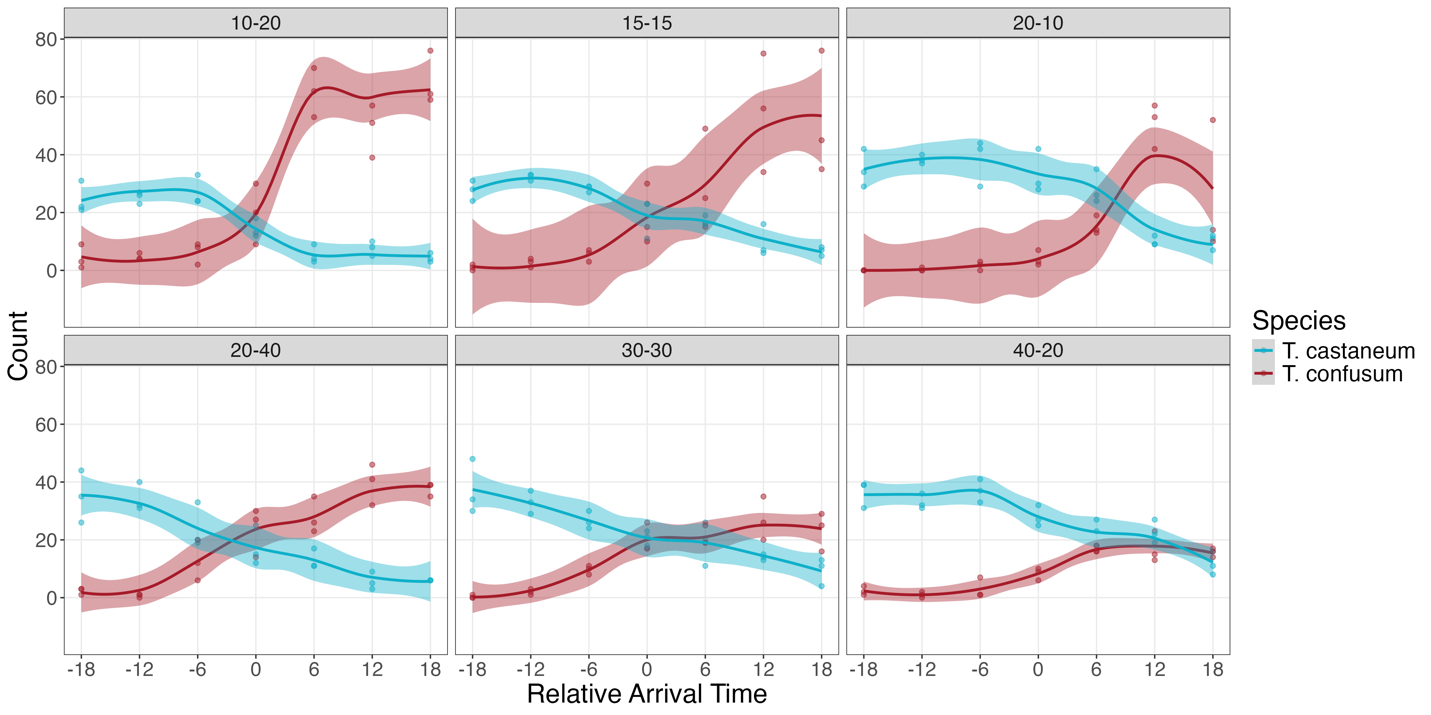

**Figure S4.** Final adult populations of vials with two generations. Numbers on top of each panel shows the initial population in the order of *T. castaneum* – *T. confusum*. Negative relative arrival times represents the early arrival of *T. castaneum*.

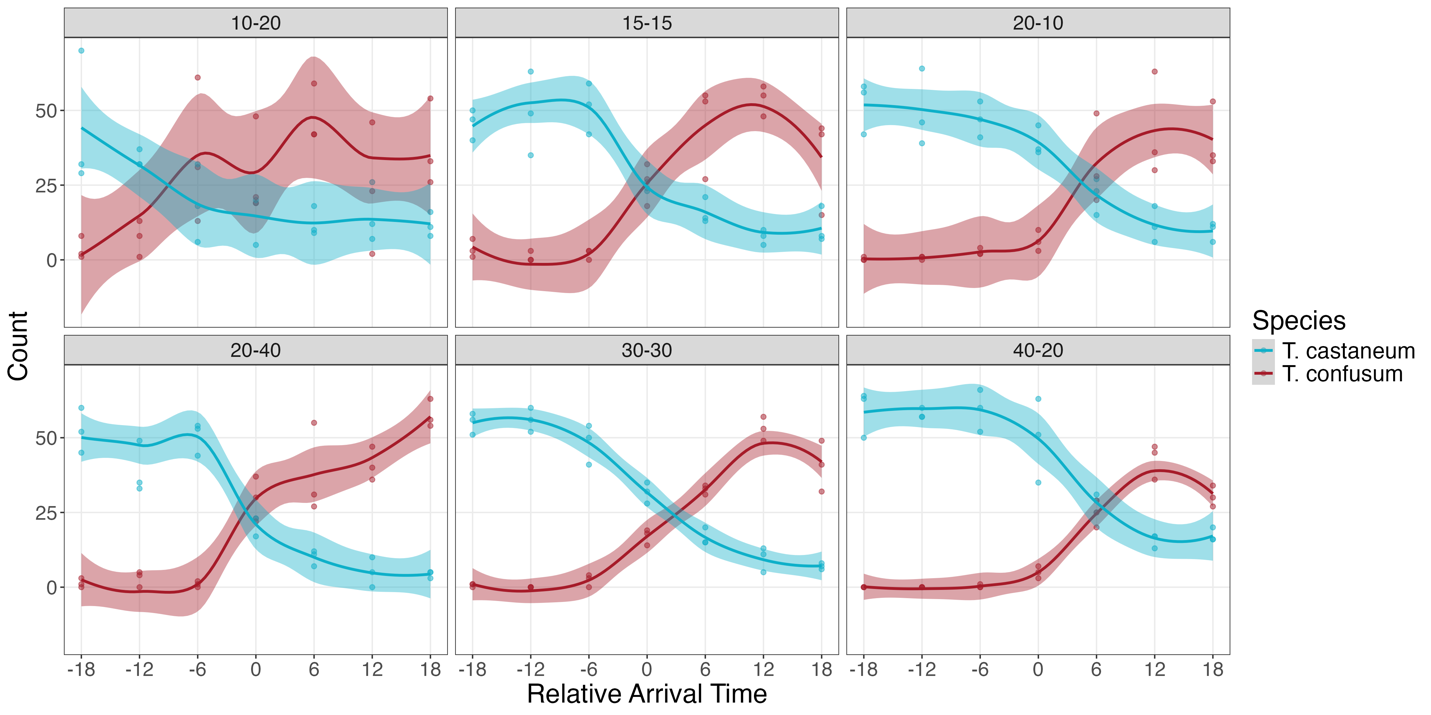

**Figure S5.** Final adult populations of vials with three generations. Numbers on top of each panel shows the initial population in the order of *T. castaneum* – *T. confusum*. Negative relative arrival times represents the early arrival of *T. castaneum*.

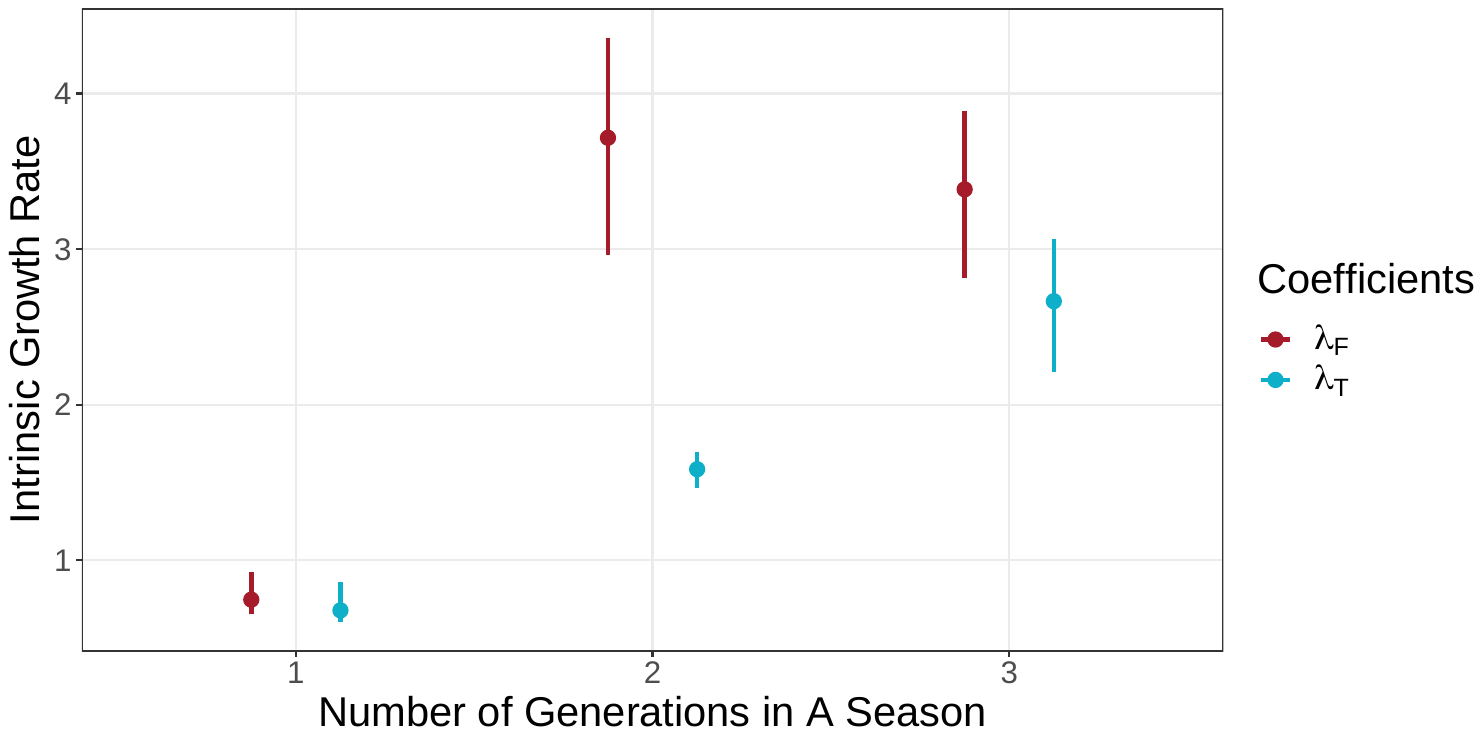

**Figure S6.** Fitted intrinsic growth rates from single-species vials. *F* represents *T. confusum*; *T* represents *T. castaneum*. Points show the medians, and line ranges show 80% credible intervals of the posterior distribution.

**Additional Experiments**

*Egg predation*

To evaluate the potential of egg predation, we added 30 eggs of one species to one vial with 8g medium containing 30 larvae of the other species at age 6, 12, and 18 days. Each vial is repeated six times. These conditions are identical to the starting conditions of the experimental vials. After one day, we sieved the vial to count number of remaining eggs. We used eggs collected less than 24 hours ago from the start of the experiment to minimize the possibility that any eggs hatched during the experiment. On average, larvae of *T. castaneum* consumed more eggs of *T. confusum* (one-sided Student’s *t* = -8.84, *p* < 0.01, n = 18): the average egg survived in vials with *T. castaneum* is 11.50 (SE = 1.40), and the average number for *T. confusum* is 25.72 (SE = 0.80). For *T. castaneum*, the consumption increases with later stages (linear regression slope of age is -0.60, *p* = 0.03, n = 18).

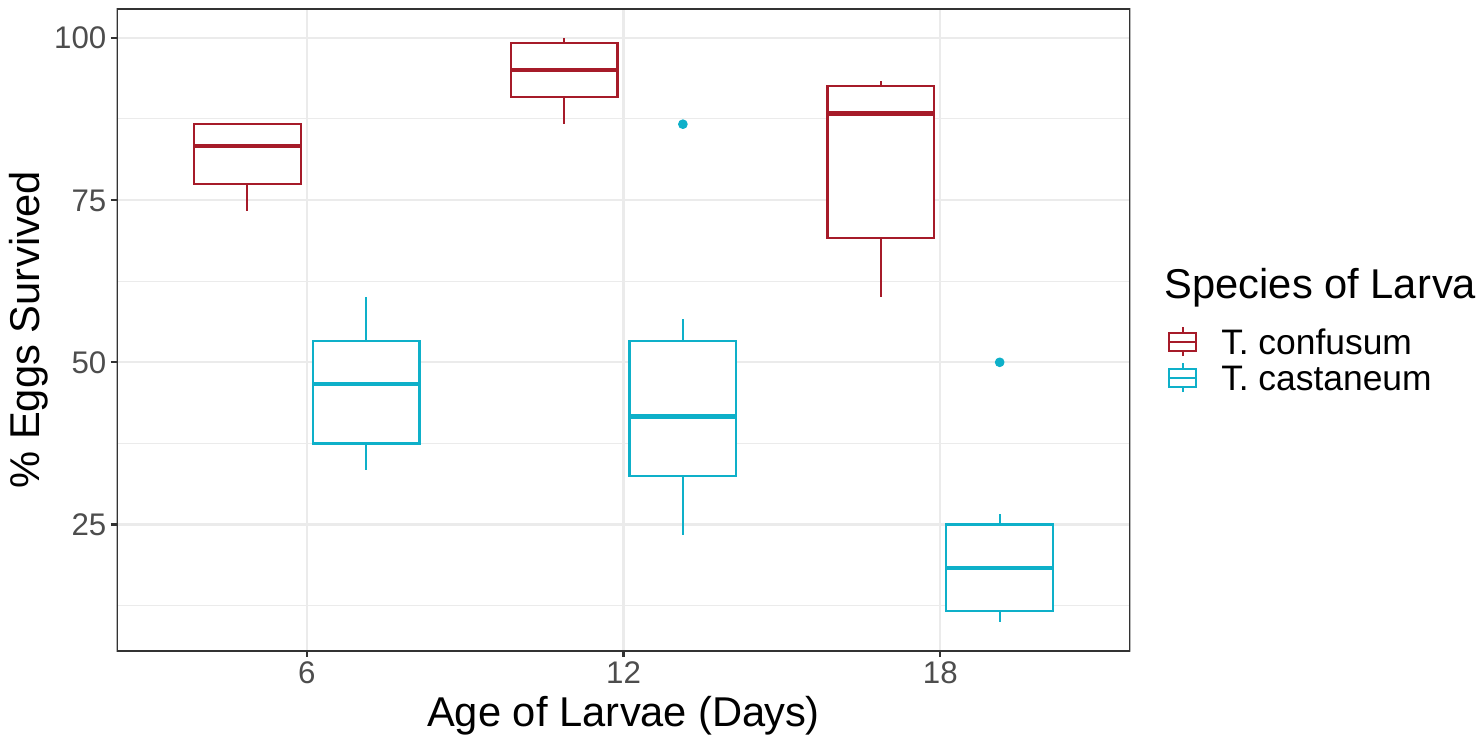

**Figure S7**. Predation of *T. castaneum* and *T. confusum* larvae on eggs of the other species.

*Fecundity*

We compared the fecundity of the two species by adding 30 randomly selected adults from our stock culture to 50g medium. We counted the number of eggs daily for five consecutive days. *T. castaneum* produced less eggs than *T. confusum* (one-sided Student’s *t* = -9.07, *p* < 0.01, n = 5), corresponding to our observations during maintenance of stock cultures and egg collection, that the latter can reach much higher abundance under lab conditions.

| **Species** | **Day 1** | **Day 2** | **Day 3** | **Day 4** | **Day 5** | **Average (SE)** |
| --- | --- | --- | --- | --- | --- | --- |
| *T. castaneum* | 72 | 102 | 107 | 103 | 99 | 96.60 (6.28) |
| *T. confusum* | 291 | 371 | 266 | 297 | 249 | 294.80 (20.92) |

**Table S4.** Number of eggs laid by 30 randomly sampled individuals over five consecutive days.
